# Rapid and fully automated blood vasculature analysis in 3D light-sheet image volumes of different organs

**DOI:** 10.1101/2022.09.14.507895

**Authors:** Philippa Spangenberg, Nina Hagemann, Anthony Squire, Nils Förster, Sascha D. Krauß, Yachao Qi, Ayan Mohamud Yusuf, Jing Wang, Anika Grüneboom, Lennart Kowitz, Sebastian Korste, Matthias Totzeck, Zülal Cibir, Ali Ata Tuz, Vikramjeet Singh, Devon Siemes, Laura Struensee, Daniel R. Engel, Peter Ludewig, Luiza Martins Nascentes Melo, Iris Helfrich, Jianxu Chen, Matthias Gunzer, Dirk M. Hermann, Axel Mosig

## Abstract

Blood vasculature represents a complex network of vessels with varying lengths and diameters that are precisely organized in space to allow proper tissue function. Light-sheet fluorescence microscopy (LSFM) is very useful to generate tomograms of tissue vasculature with high spatial accuracy. Yet, quantitative LSFM analysis is still cumbersome and available methods are restricted to single organs and advanced computing hardware. Here, we introduce VesselExpress, an automated software that reliably analyzes six characteristic vascular network parameters including vessel diameter in LSFM data on average computing hardware. VesselExpress is ~100 times faster than other existing vessel analysis tools, requires no user interaction, integrates batch processing, and parallelization. Employing an innovative dual Frangi filter approach we show that obesity induces a large-scale modulation of brain vasculature in mice and that seven other major organs differ strongly in their 3D vascular makeup. Hence, VesselExpress transforms LSFM from an observational to an analytical working tool.

## Introduction

Vascular reorganization is a key process accompanying various pathophysiological conditions. For instance, sterile tissue inflammation induced by hypoxia-ischemia in stroke, myocardial infarction or tumor development leads to massive vascular remodeling that critically determines tissue fate^1–3^. Due to its unprecedented 3D visualization capacity light-sheet fluorescence microscopy (LSFM), which has already been employed for analyzing a variety of organs^2,4–7^, also allows a thorough exploration of disease-associated vessel reorganization. The labeling of blood vessels can be achieved by the injection of fluorescently labeled endothelial-specific antibodies^4^, the perfusion of animals with fluorescent dyes^8^ or the use of transgenic animals in which fluorescent reporter proteins are expressed under endothelial-specific promoters^9^. After vessel labeling, vertebrate organs need to be optically cleared before LSFM^10–12^. The power of LSFM enables high-resolution tomograms of whole cleared organs to be acquired quickly. The processing and analysis of the large (10-100 GB) data sets, however, can be laborious, which strongly inhibits the application of LSFM for large-scale quantitative studies of vessel systems.

Even though methods have been put forward that allow the quantification of blood vessel structures, even in large organs such as the murine brain^6,7^, their application has only been demonstrated in the investigated organ, hence lacking general usability. Furthermore, they lack processing scalability, depend on advanced computing hardware, and even with advanced hardware, they are too specific to allow for the systematic analysis of more complex pathophysiological settings by non-experts. Recently, efforts have been made to make 3D vessel analysis accessible for non-experts^29^. However, by requiring pre-segmented data, the tool cannot be used out-of-the box with raw microscopic data and thus fails to provide a complete workflow. Furthermore, other existing solutions are neither computing platform independent nor do they provide parallelizable workflow processing for medium or high throughput studies. The non-modular software design of existing solutions also lacks flexible adjustment of individual processing steps. In short, the lack of reliable and robust methods with modular and parallelizable workflow management for the quantification of phenotypic vessel features from the large 3D image stacks is a major limiting factor. Overcoming this limitation promises a novel paradigm to study disease processes and, potentially, also screening compound libraries for effects on whole organ vascularization^13^.

Here we present VesselExpress, a software to fully automatically analyze LSFM 3D data of blood vessel systems. It allows fast and reliable image analysis including image processing methods, graph construction and analysis, which are modularly assembled in a workflow. VesselExpress enables high-volume analyses, supported by corresponding algorithms, computational tools^14,15^, and workflow management systems^16^. With VesselExpress we analyzed blood vessels from different murine organs which were labeled either with fluorescein isothiocyanate (FITC)-albumin or fluorescent antibodies against CD31. Using an innovative strategy that combines statistics-based thresholding with two customized Frangi filter segmentations, we were able to extract a comprehensive set of microvascular network characteristics that includes reliable information on microvascular length, branching and diameter. Obtained measurements are well in line with conventional microscopic quantifications and match or exceed the performance of semi-automated analyses, while operating orders of magnitude faster and with far less hands-on time. Furthermore, VesselExpress is ready-to-use through a browser-based web interface without expert knowledge, freely available and platform-independent.

## Results

### VesselExpress software and workflow

VesselExpress is open-source software designed for rapid, fully automated and scalable analysis of blood vessel trees in 3D data sets from LSFM sequences. It is developed in Python and freely available (https://github.com/RUB-Bioinf/VesselExpress). VesselExpress processes raw microscopic images of blood vessels in parallel and outputs quantified phenotypical data without user interaction, fully automatically. The software is deployed as a Docker^14^ container which includes the necessary run environment and can be executed from all major operating systems via command line or the web interface integrated into the Docker image.

The workflow is illustrated in Figure 1. VesselExpress accepts raw LSFM 3D images in the standard image file formats as input. Blood vessel staining and tissue clearing (Fig. 1A) are required as sample preparation steps for the subsequent imaging of blood vessels in high resolution (Fig. 1B). The raw image volumes (Fig. 1C) are then processed fully automatically without any user interaction (Fig. 1D). The processing steps include segmentation (Fig. 1D1), skeletonization (Fig. 1D2) and graph construction with analysis (Fig. 1D3). In the first step, the raw images are segmented using workflows extended from Allen Cell and Structure Segmenter^17^. The same workflow is employed but with different parameters optimized for different organs. Users can choose to load our preset parameters for a specific organ or further fine-tune the parameters within a Napari^18^ plugin. In the second step, the vessels’ centerlines are extracted from the binary images through the parallel thinning algorithm^19^ implemented in the scikit-image Python package^15^. The centerlines are then transformed into undirected graphs by using the Python 3scan toolkit^20^. Finally, the vessels are traced in a depth-first search (DFS) and nine phenotypic features (Fig. 1E) are calculated. These steps are automated in a pipeline integrated into the workflow management system Snakemake^16^ which enables high scalability so that the run time of the software automatically benefits from larger RAM and more CPU cores without having to make any changes to the code. This allows comprehensive studies to be carried out within a short time. Due to the modular software design, each processing step can also be executed individually. Therefore, each function (“module”) can be easily exchanged with custom functions, if preferred, while maintaining the integrity of the pipeline. The pipeline outputs the nine quantified phenotypical features as text files which can be directly statistically analyzed and visualized as required. Furthermore, 3D TIF images of the segmented and skeletonized vasculature as well as images of the graph, branching and terminal points are provided. Optionally the 3D vessel tree can automatically be rendered in Blender^21^. Images of the rendered vessel tree can be provided along with the corresponding Blender project file that may be used for visual inspection and presentation.

**Fig. 1.**
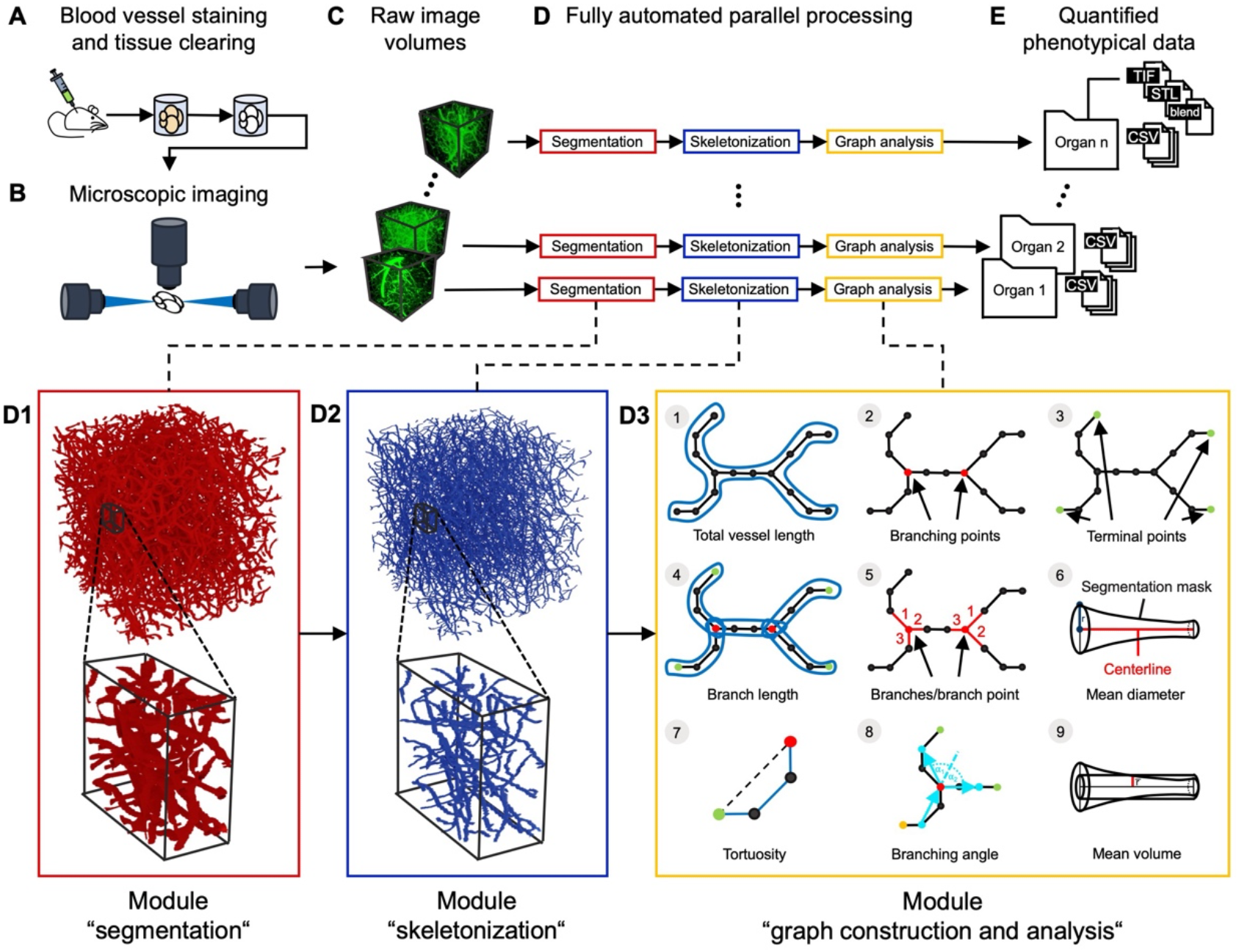
Pipeline for automatic quantitative blood vessel analysis. **(A)** Sample preparation includes blood vessel staining and tissue clearing. **(B)** Microscopic imaging of blood vessels. Generated blood vessel images **(C)** are processed via VesselExpress **(D)**. In the segmentation the raw images are binarized into foreground, representing the vessels, and background **(D1)**. The segmentation mask is used to extract the vessels’ centerlines **(D2)**. Undirected graphs are constructed from the skeleton mask and used for subsequent data analysis **(D3)**. The results are quantified phenotypical data tables for statistical analysis and image masks for visual inspection of segmentation and skeletonization **(E)**.

### Benchmarking and validation

We generated high-resolution LSFM data sets of mouse brains with their blood vessels labeled by injection of FITC albumin (Fig. 2A, B) to benchmark VesselExpress against Imaris, a widely used software tool for 3D vessel analysis. When these data sets were analyzed using VesselExpress on a workstation or multi-CPU server with large amounts of RAM, we observed a linear increase in computing time that was proportional to the ROIs volume (Fig. 2C). In example images from mouse brains, ROIs with a size of up to 0.25 mm^3^, i.e., 250 MB’s of 16-bit intensity images, could be analyzed in parallel on an office PC. It is worth mentioning that, unlike other comparable analysis methods, the performance of VesselExpress depended on the availability of RAM and the number of CPU cores. Furthermore, our code uses the Dask^22^ package internally to overcome limitations where the data set is larger than the available RAM on a machine. Interestingly, the vessel length density (Fig. 2D) and total length (Fig. 2E) vs ROI volume showed only a slight deviation from constancy or linear growth respectively, with larger ROIs likely representing the highly heterogeneous capillary density in murine brains that becomes obvious at deeper regions of e.g. the cortex^7^. A head-to-head comparison of 12 or 48 different ROIs of the same brain regions showed that VesselExpress is about 70 or 95 times faster, respectively, compared to our reference system based on Imaris (Fig. 2F). This is due to the capability for parallelized batch processing of arbitrarily large ROI sets in VesselExpress, while Imaris requires each ROI to be annotated manually and separately. Nevertheless, the quantifications obtained from both approaches differed only insignificantly (Fig. 2G). We benchmarked the scalability of VesselExpress by investigating run times on an increasing number of 508 × 508 × 1000 μm ROIs. Our experiments involved 1024 ROIs on a workstation and high-performance computing hardware, respectively. Supplemental Figure 1 confirms that VesselExpress scales linearly with respect to input size and concurrency.

**Fig. 2.**
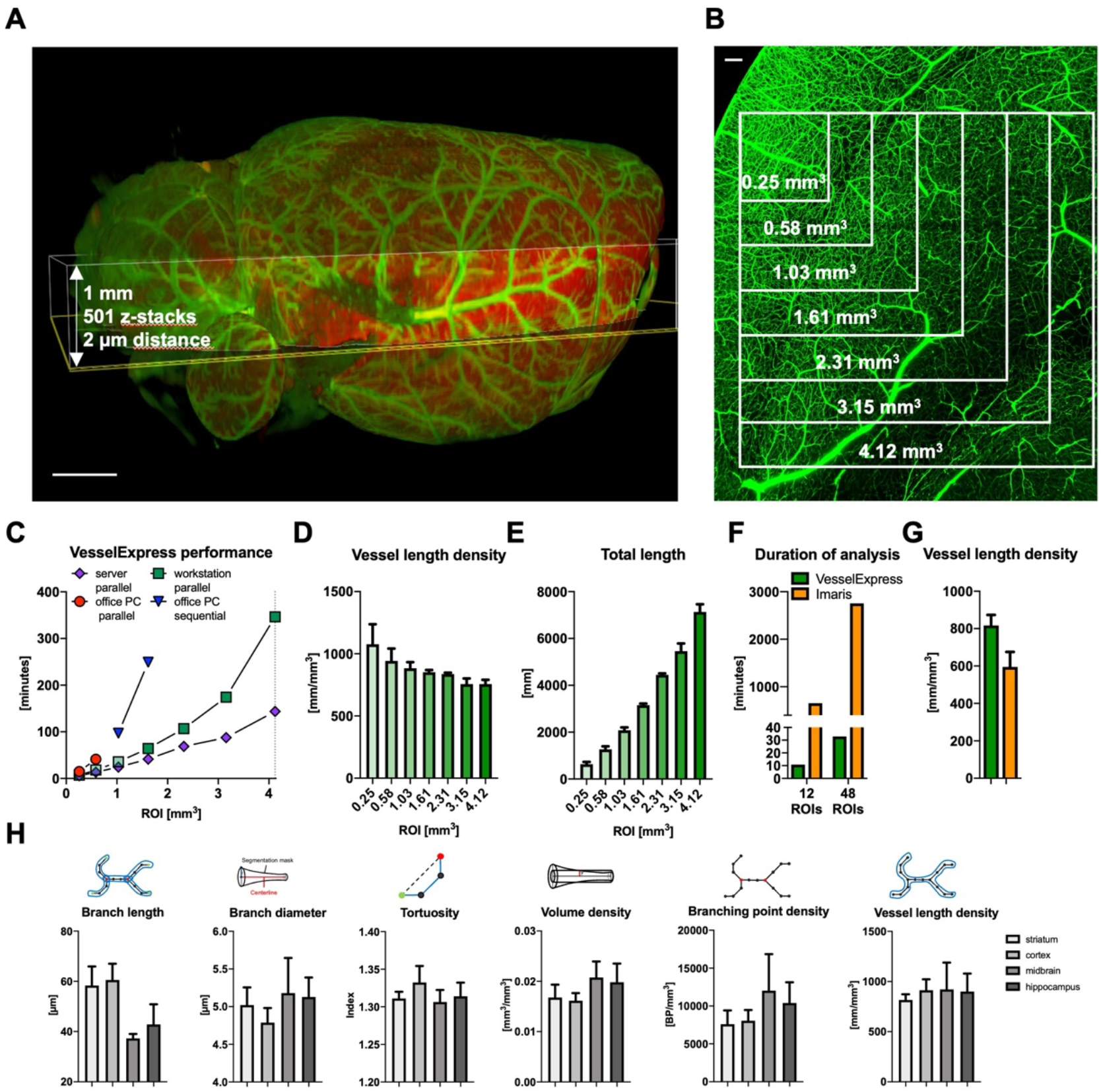
Analysis of FITC-Albumin labeled brain vessels using automated pipeline VesselExpress. **(A)** Overview of brain vasculature in a mouse brain scanned with LSFM. Highlighted area represents region in which detail scans for vessel analyses in **(C-G)** were performed. **(B)** Maximum intensity projection of detailed image from LSFM scan using 6.64 × magnification and a step size of 2 μm. Inlets showing the region of interest used for vascular analysis in **(C-E)**. **(C)** Duration of vascular analysis in VesselExpress using four ROIs each in parallel with increasing image sizes. Analysis was performed on different devices. Using an office PC containing an Intel Core Duo i9 processor with 2.40 GHz, 8 cores and 32 GB RAM, parallel analysis was possible with ROIs of a maximum size of 0.58 mm^3^ (red circles). Using one ROI per run, it was possible to analyze a ROI with 2.31 mm^3^ and total run times were summed up (blue triangle). Green squares represent a workstation equipped with an Intel Xeon W-1255 CPU with 3.30 GHz and 512 GB RAM whereas violet diamonds display runtime of the very same ROIs on a server working with an Intel Xeon E7-8890 v4 CPU with 2.20GHz, 96 cores and 1.97 TB RAM. Results from this analysis are displayed as vessel length density **(D)** and total vessel length **(E)**. Duration **(F)** and Vessel length density **(G)** from the analysis of either 12 ROIs or 48 ROIs with an image size of 0.25 mm^3^ using the automated pipeline VesselExpress or sequentially analysis using Imaris software (Bitplane, Oxford Instruments). **(H)** Analysis of FITC-Albumin labeled vascular structure performed in 4-12 ROIs of striatum, cortex, midbrain, hippocampus and corpus callosum of brains from up to 6 male mice using VesselExpress. Scale bars represent 1 mm in (A) and 100 μm in (B).

We investigated the validity of VesselExpress outputs along several lines. The first evidence of the validity was provided by the insignificant deviation between VesselExpress and the Imaris-based workflow (Fig. 2G). Since Imaris requires inspection and adjustment of intermediate results by the user, it can be considered a gold standard reference. The agreement between the semi-automated Imaris workflow and our highly automated VesselExpress was also supported by detailed visual overlays of specific branching points (Supplemental Figure 2). As further validation, we generated synthetic data that model blood vessels with known lengths and diameters as ground truth. As shown in Supplemental Table 2, vessel lengths determined by VesselExpress deviated by less than 2%, while the average vessel diameter deviation was below 15%. We analyzed six typically used vesselness features provided by VesselExpress in different regions of the brain (Fig. 2H) and found distinct values for every brain region that are well in line with published data^6,7^. We observed that obtaining vessel diameters in line with published literature crucially relied on adequate image segmentation as the first analysis step. Our vessel-specific Frangi-filter based apporoach (see supplemental methods) yielded vessel diameters that very well matched with data from histological analyses, summing up to 1-2% of fractional brain volume density (Fig. 2H)^23^.

Comparing VesselExpress with the semi-automated Imaris-based segmentation and the recently published VesselVio^24^ using the segmented data from VesselExpress, produced highly similar results for the synthetic data (Supplemental Table 2), whereas vessel length density in real brain data was ~18% higher in VesselExpress than in VesselVio and Imaris (Supplementary Figure 3A). This effect disappeared after vessel smoothing which introduces an artificial shortening of vessel lengths, so that we refrained from using it in further analyses. Smoothing is, however, implemented as an optional choice in VesselExpress. Importantly, VesselVio does not support the crucial step of image segmentation, so that the comparison required input images that were pre-segmented using VesselExpress. Furthermore, VesselVio is much slower than VesselExpress and is limited in terms of automated analysis (Supplementary Figure 3B).

### Obese mice show defects in brain vascularization

To demonstrate that VesselExpress enables automated analysis of data sets addressing a specific scientific question, we fluorescently labeled vessels in male 9-12-week-old C57BL/6 wild-type or homozygously leptin-deficient (ob/ob) mice of the same age by perfusion with FITC-albumin containing hydrogel. A point mutation of the leptin gene in these animals leads to the development of hyperlipidemia and associated comorbidities such as hyperglycemia, hyperinsulinemia and infertility^25^. We then evaluated ROIs at different rostrocaudal levels of the brain including striatum, cortex, midbrain and hippocampus using VesselExpress (Fig. 3). Our structural data revealed a substantial impairment of microvascular network integrity in different brain regions of ob/ob compared to wildtype mice (Fig. 3A-G). In striatum and cortex of obese mice, we observed significantly lower branch lengths, branch diameters, and tortuosity indices resulting in lower vessel volumes compared to wildtype mice, while the total vessel length density was indistinguishable at all sites. A more detailed analysis of vessel length density in relation to small (<4 μm), medium (4 to 6 μm) and large diameters (>6 μm; Fig. 3H) revealed that obese mice have more thin vessels (<4 μm) but fewer vessels larger than 4 μm (Fig. 3I) in the striatum. Hence, LSFM followed by automated VesselExpress analysis can uncover subtle vascular changes in the brain associated with a metabolic disease. Interestingly, mice with a defect in a related protein, the leptin receptor (db/db), which are characterized by obesity and diabetes, were recently shown to reveal a biphasic vascular development with a juvenile hypovascularization followed by aberrant hypervascularization in later adulthood^26^, although reported densities for branch points in that earlier study assessed by LSFM were lower than our data (Fig. 3A) and data reported elsewhere in the literature^7^. To the best of our knowledge, the consequences of leptin deficiency for brain microvascular networks had not been studied by LSFM. Obesity and diabetes evoke multifaceted vascular changes in the brain^27^. Obesity triggered by a high-fat diet induces a significant increase in vascular density, especially in the hippocampus under physiological conditions as assessed by LSFM^28^, but compromises cerebral microvascular proliferation and remodeling post-ischemia^29^. Importantly, it has been shown in humans that obesity negatively influences brain perfusion^30^. Hence, LSFM data in mice appear to show the microvascular correlate of this finding.

**Fig. 3.**
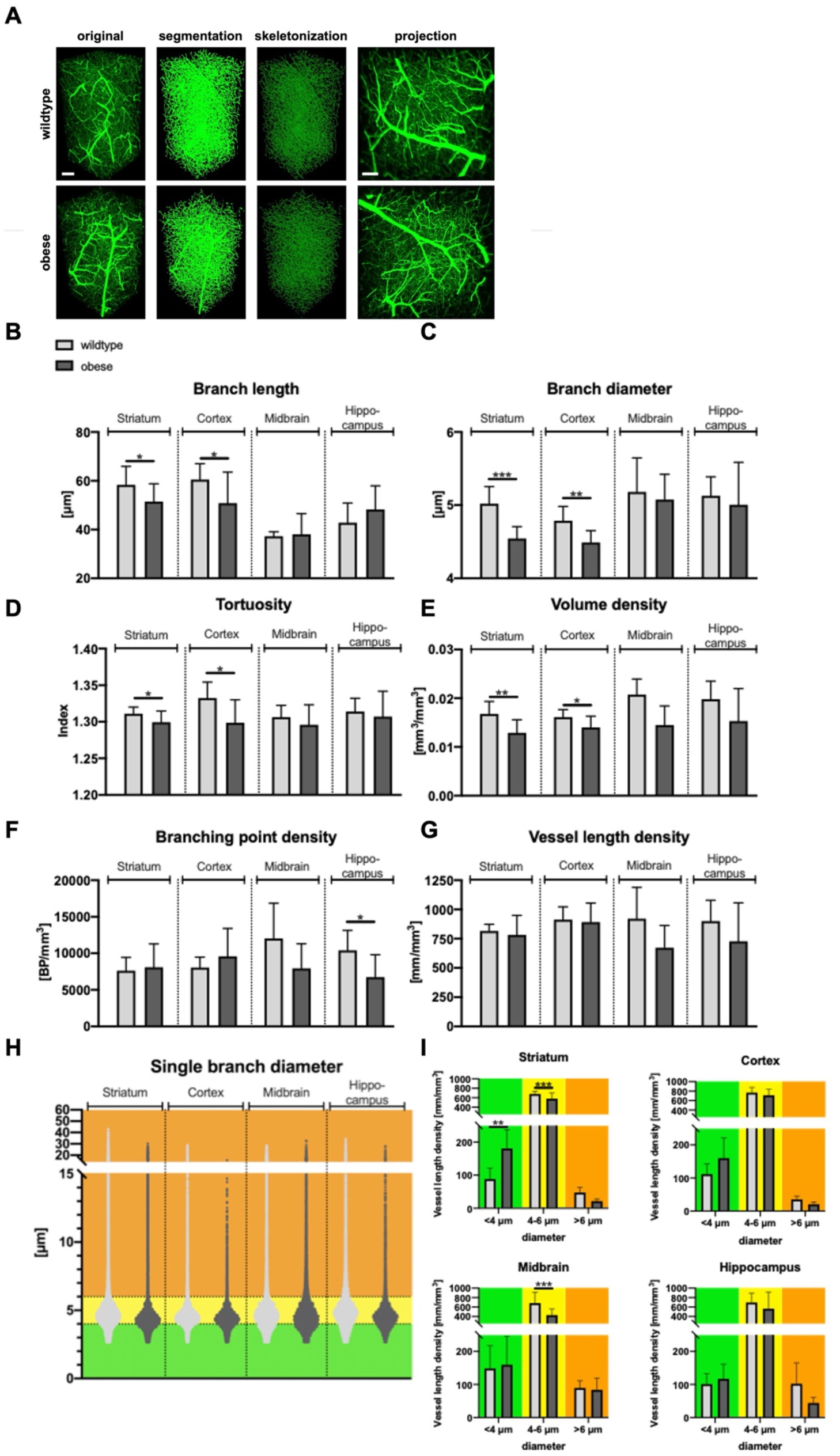
Vascular changes in brains of wildtype compared to obese mice. **(A)** Representative images of striatal brain regions showing a 3D view of original data, segmented pictures and skeletonized images provided by VesselExpress as well as a maximum projection of stacked image (right). **(B)** VesselExpress analysis of brain vessels reveals significant decrease in branch length, branch diameter, Tortuosity and branch volume density in the striatum and cortex in obese mice compared to age-matched wildtype mice. **(C)** dot plot displaying single vessels according to diameter in different brain regions in wildtype (light grey) and obese mice (dark grey). Dashed lines represent thresholds used for the analysis of vessel length density in **(D). (D)** Vessel length density in vessels with diameters <4 μm, 4-6 μm or larger than 6 μm shows increased Vessel length density of obese mice in smaller vessels and reduced vessel length density in larger vessels in the striatum. Data are means ± SD values. *p≤0.05/**p≤0.01/***p≤0.001 compared with wildtype; n = 4–12 ROIs per group; analyzed by students T-test (in B) or two-way Anova using Sidaks multiple comparison test in (D). Scale bars represent 100 μm.

### VesselExpress can analyze the vasculature of many organs

An obvious key requirement for a general-purpose vessel analysis tool is the transferability across different organs. Currently available tools, while certainly extremely powerful for their specific use in brain^6,7^, are not able to analyze LSFM data sets of other organs. Hence, we validated VesselExpress for analyzing blood vessels of seven different organs and an additional heart data set from a different series. We prepared six animals by injection of FITC-albumin, isolated the organs and generated high-resolution data sets of blood vessels (Fig. 4A). We found that VesselExpress could effectively extract complex vessel features from each organ. This analysis showed very similar values for vessel tortuosity, while all other features varied strongly between organs, thereby showing varying degrees of value spreading or homogeneity between samples. The liver turned out to be the most heavily vascularized organ with the highest vessel length and volume density (Fig. 4B). Interestingly, close observation of the vessels by diameter revealed a different distribution depending on the organ. Thus, a predominant number of the vessels in the tongue and heart were thinner than in other organs, while vessels of the liver, muscle and ear were thicker than those in the brain (Supplemental Figure 4A). The distribution of the length of single vessels is comparable throughout the different organs, although brain and muscle reveal a bigger proportion of long vessels (Supplemental Figure 4B). This demonstrates that LSFM-datasets from various organs are ready-to-use when applying the provided organ-specific configurations in the pipeline without the need for further modifications. To test if VesselExpress could also be used for studying vascular features in animals exhibiting antibody-mediated labeling of endothelial cells, we finally examined microvascular network characteristics in murine hearts exposed to myocardial ischemia/ reperfusion (I/R) injury, to which we intravenously delivered AF647-labeled anti-CD31 antibodies prior to animal sacrifice^2^. Our study showed that VesselExpress was able to image microvessels in these samples with results very similar to previously published data^2^ (Supplemental Fig. 5; Fig. 4A,B). Furthermore, VesselExpress also analyzed brain vessels labeled simultaneously using FITC-albumin hydrogel as well as AF647-CD31 antibody with very comparable results (Supplemental Fig. 6). In addition, VesselExpress is not restricted to LSFM images. As shown in Supplemental Fig. 7, results from VesselExpress in brain vasculature imaged with confocal microscopy yielded similar results regarding branch length, branch diameter, tortuosity, branch volume and branching point density.

**Fig. 4.**
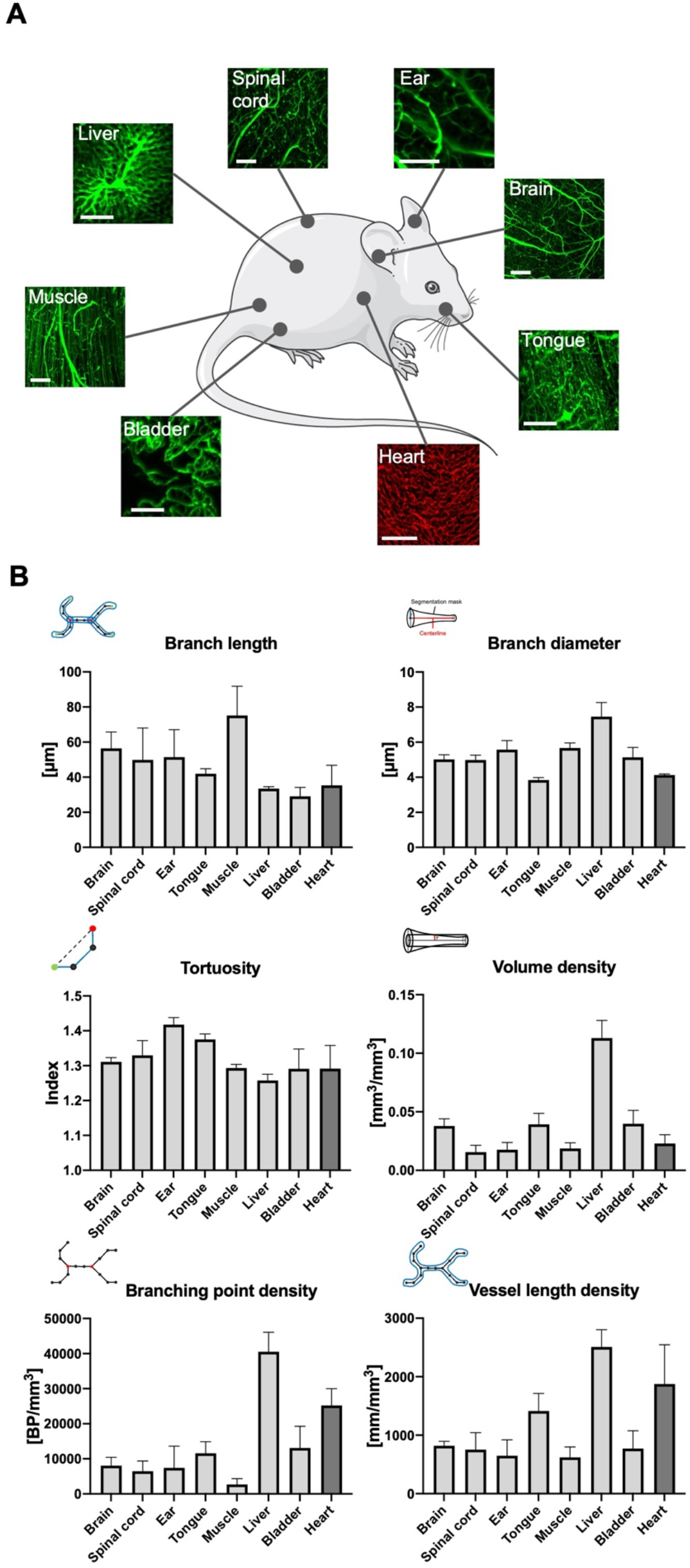
VesselExpress is applicable for a variety of organs of mice. A total of 6 mice were perfused with FITC-albumin gelatin hydrogel and indicated organs **(A)** were removed, dehydrated, and cleared. In a proof of evidence experiment, the heart was labeled separately by injection of AlexaFluor-647-labeled anti-CD31 into additional mice. LSFM was performed as described before and vascular parameters including **(B)** branch length, branch diameter, tortuosity, branch volume density, branching point density and vessel length density were calculated. Scale bars represent 100 μm.

## Discussion

With the development of advanced 3D imaging methods and LSFM in particular, there is a high demand for automated analysis of large data sets. Existing software is able to analyze individual 2D or 3D images^1,31^, but currently the analysis especially of 3D data is disproportionately time-consuming and requires frequent and extensive hands-on time. A major bottleneck in existing solutions is the first and essential processing step of segmentation, which critically affects analysis outcome and thus often involves hands-on intervention. Our implementation of VesselExpress in a script-based workflow management system facilitates a paradigm shift in the application of LSFM for comprehensive measurement campaigns since the analysis of the ensuing data is removed as a bottleneck. The workflow-based approach systematically automates data analysis, following high-throughput paradigms of high-content image analysis approaches. At the same time, a user-friendly environment allows initial installation and execution of the code by non-experts. Furthermore, the software design reduces the computational time by up to two orders of magnitude, thus making higher throughput studies and analyses possible on broadly available commodity hardware. With these considerations in mind, VesselExpress represents a fully automated analysis method for LSFM datasets with low barrier to entry. It is easily accessible via the provided Docker image operated by a simple command line or via an integrated web interface with few required configuration steps and does not need technically sophisticated hardware, specific operating systems or advanced computer skills compared to other methods, while at the same time producing comparable results^6,7^. 16-bit images with a size of 250 MB can easily be analyzed in parallel on a conventional office PC within 3 minutes per ROI compared to average 54 minutes for the analysis of the same image in Imaris. File sizes of 250 MB represents ROIs with a volume of 0.25 mm^3^ for light-sheet images scanned with optimized resolution in 16-bit depth. For the analysis of larger images or larger numbers of images, however, a computer with a powerful CPU and a large-sized RAM is recommended to reduce the processing time with the parallel pipeline. In terms of processing time this facilitates analyses in VesselExpress that are impractical in comparable pipeline-based algorithms which only enable sequential processing and do not include all required analysis steps^7,24^. We were also able to show that parallel processing allows thousands of images to be processed in batches on high-performance computing hardware.

A downside of existing microvascular segmentation solutions is that calculated vessel diameters overestimate true microvessel diameters, resulting in, depending on the mode of image acquisition, image axis and data processing, mean brain capillary diameters >5 μm^1^ or even significantly >10 μm^6^. In confocal microscopic analyses, the diameter of capillary lumina in the cerebral cortex of C57Bl/6 mice and Wistar rats was shown to be 3.5-4.0 μm and 5.4±1.5 μm, respectively^23,32^, whereas that in the rat cerebral cortex was 4.2±1.2 μm in Indian ink-stained corrosion casts^33^. As a consequence of vessel diameter overestimations, the majority of existing LSFM studies including our studies refrained from statistically evaluating microvascular diameters^28,34–36^. Here we now made particular efforts to obtain reliable vessel diameters using an innovative strategy that combines statistics-based thresholding with up to two customized Frangi filter segmentations (see supplemental methods). By this approach we computed mean microvessel diameters in the cerebral cortex of 4.8±0.2 μm, summing up for the whole brain to 1-2% of fractional volume density, which very well matches data from conventional histological analyses^23^. Hence, we can extract comprehensive sets of network characteristics that comprise information on microvessel length, branching and diameters. With this tool, we for the first time detected statistically significant differences in the vascular make-up of leptin-deficient ob/ob compared with wild type mice, reflected by an increase of thin (<4 μm) microvessels. At present, VesselExpress analyses are optimized for vessels filled with a hydrogel containing a fluorescent dye. Labeling of vessels using endothelial-specific antibodies poses particular challenges, since endothelial labeling results in hollow tubes. In case of the brain and heart vascular analysis was similarly possible following endothelial labeling with anti-CD31 antibodies. We so far did not evaluate the utility of VesselExpress for studying CD31-labelled vascular networks in tissues with larger caliber vessels. Future studies will examine which pipeline adaptations are required to enable robust network analysis in such tissues. The modular design of VesselExpress would, however, allow to easily expand the pipeline for required additional components.

Taken together, our approach enables the transition of LSFM from observational single-case studies to high content analyses that systematically and quantitatively evaluate organs and tissues. Our results demonstrate that LSFM can generate detailed vessel tomograms of multiple organs and thus could form the basis for a novel type of organ map that considers the 3D organization of higher-order structures. Since blood vessel densities and characteristics are important e.g. for the efficiency of chemotherapy of solid cancers^37^, VesselExpress also bears substantial potential for broad applicability^38^.

## Materials and Methods

### Animals

Animal experiments were performed in accordance with the regulations of the National Institute of Health Guidelines for the Care and Use of Laboratory Animals in compliance with ARRIVE guidelines and the permission of local authorities (Ministry for Environment, Agriculture, Conservation and Consumer Protection (MULNV) of the State of North Rhine-Westphalia). Organs of interest were removed in Ketamin/Xylazin anesthesia from C57BL/6J mice after perfusion with 40 ml 1 × PBS followed by 40 ml 4% PFA and 10 ml FITC-albumin containing gelatine hydrogel as described before^1^. For analysis of hearts, mice were subjected to a myocardial (I/R) injury in vivo, as reported^2^. For this purpose, the animals’ rib cage was opened through a left lateral thoracotomy via which the left coronary artery (LCA) was ligated. After 45 min of ischemia, reperfusion was allowed for 5 days. For labeling of endothelial cells, these mice received 5 μg Alexa 647-coupled anti-CD31 antibody (Cat No. 102516, BioLegend) i.v. in a total volume of 150 μl PBS 10 min prior to sacrifice by cervical dislocation followed by animal perfusion with PBS^2^. All tissues used in this study were dehydrated with increasing concentrations of tetrahydrofluran (THF) for 12 hours each and cleared using ethylcinnamate (ECi), as outlined before^1^.

### Light sheet data acquisition

Cleared organs were imaged using a light sheet UltraMicroscope Blaze™ (LaVision BioTec (Miltenyi), Bielefeld, Germany) equipped with a supercontinuum white light laser source for bidirectional light sheet illumination, an sCMOS camera having a 2,048 × 2,048 chip of 6.5 μm pixel size, and three dipping objectives with magnification 1.1X (0.1NA), 4X (0.35NA) and 12X (0.53NA), which could be combined with additional tube lens magnification of 0.6×, 1×, 1.67× or 2.5×. Serial optical imaging was performed by exciting the FITC-albumin, tissue autofluorescence or AF647-anti-CD31 labeled vessels using excitation-emission bandpass filter combinations of 470/30-525/50, 560/40:620/60, and 630/30:680/30 respectively. Overview images were acquired at 1.1× magnification with 10 μm steps in the axial direction. Detailed images of blood vessels were acquired at 6.68× magnification with 2 μm steps in the axial direction using the dynamic focus of the light sheet illumination at the highest NA for optimal axial resolution.

### VesselExpress analysis

Vascular analysis using the automated pipeline VesselExpress was performed in organs derived from the same group of mice perfused with FITC-albumin after transcardial perfusion with PBS and 4% PFA as described above. A detailed description of the Snakemake workflow, segmentation, skeletonization, graph construction and analysis, rendering and validation is included in the supplementary methods. For a detailed analysis of vessels, image stacks acquired with 6.64× magnification and a step size of 2 μm in ventrodorsal direction were used. From each of these image stacks, ROIs covering representative vascular structures of each tissue were obtained. The size of ROIs is listed in Supplemental Table S1. Files with a size of up to 3 or 4 GB in 16-bit TIF format were analyzed in parallel on a workstation containing an Intel® Xeon® W-1255 CPU with 3.30 GHz, 10 cores and 512 GB RAM or on a server equipped with an Intel® Xeon ® CPU E7-8890 v4 with 2.20GHz, 96 cores and 1.97 TB RAM, respectively. Images with a file size of up to 1.5 GB were analyzed in parallel or sequentially on an office PC containing an Intel® Core i9 CPU with 2.40 GHz, 8 cores and 32 GB RAM.

### Vascular quantification using Imaris

For detailed vascular quantification image stacks acquired with 6.68× magnification and a step size of 2 μm in ventrodorsal direction were also used. From each of these image stacks, ROI covering representative vascular structures of each tissue. In these ROIs, microvasculature was analyzed after network modelling using the Imaris 3D rendering software filament tracer tool. The very same ROIs were used for VesselExpress analysis.

### Statistics

Data are expressed as mean ± standard deviation (SD). In case of comparisons between 2 groups, two-tailed t tests were used. P values ≤0.05 were defined to indicate statistical significance. The statistical details are given in the figure legends. Statistical analyzes were performed using GraphPad Prism version 7.0 software.

### Source code and data availability

The source code is publicly available from https://github.com/RUB-Bioinf/VesselExpress. The Napari plugin for segmentation parameter tuning is available from https://github.com/MMV-Lab/vessel-express-napari and can also be found on the Napari Hub (https://www.napari-hub.org/). Example data is available online via Zenodo: https://doi.org/10.5281/zenodo.5733150

## Funding

This study was supported by the Deutsche Forschungsgemeinschaft () to DMH and MG.

## Author contributions

Conceptualization: JC, MG, DMM, AM

Methodology: PS, NH, AS, SDK, JC, MG, AM

Software: PS, NF, SDK, DS, LS, JC, AM

Validation: PS, AS, JC, MG, DMM, AM

Formal Analysis: PS, NH, AS, JC, MG, DMM, AM

Investigation: PS, NH, AS, SDK, YQ, AMY, JW, LK, SK, ZC, AAT, VS, AAT, LMNM, IH, JC

Resources: PS, AS, SDK, YQ, AMY, JW, AG, MT, DRE, PL, LMNM, IH, MG, DMM

Data Curation: PS, NH, AS, SDK, DS

Visualization: PS, NH, NF, SDK, DS, JC

Supervision: JC, MG, DMM, AM

Writing—original draft: PS, NH, JC, MG, DMM, AM

Writing—review & editing: PS, NH, AS, SDK, VS, JC, MG, DMM, AM

## Competing interests

Authors declare that they have no competing interests.

## Data and materials availability

VesselExpress source code is available from https://github.com/RUB-Bioinf/VesselExpress. Data are publicly available from https://zenodo.org/record/6025935#.YvtjUi8RqJ8.

## Supplementary Materials for

**Fig. S1:**
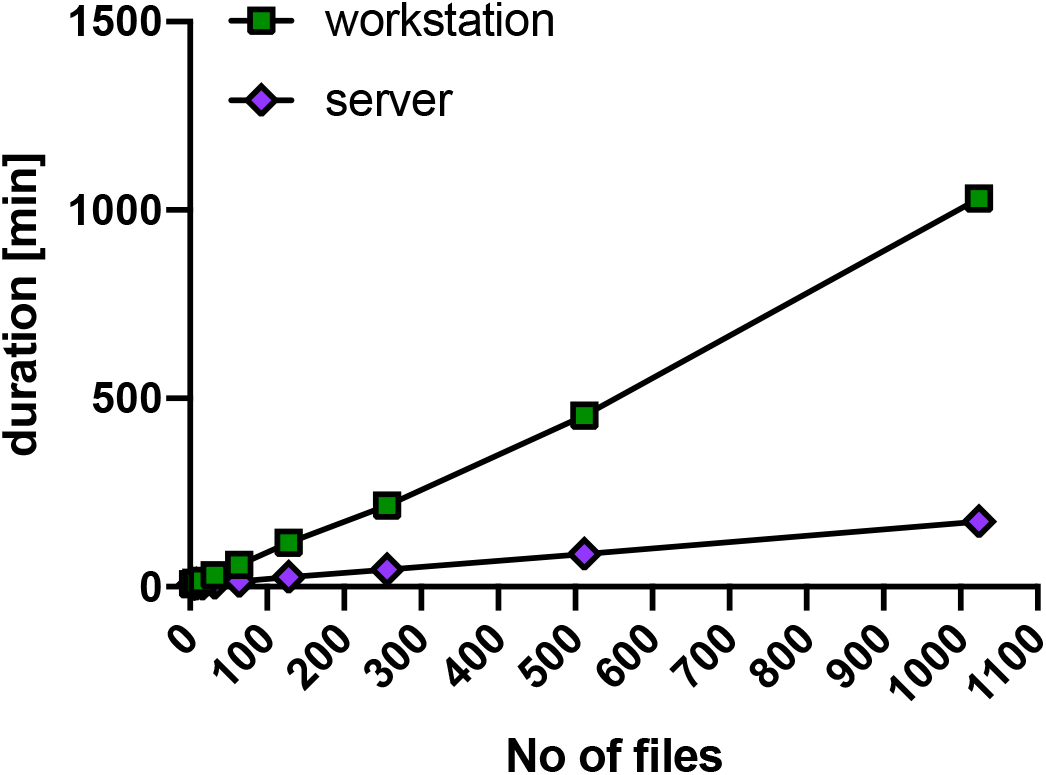
Duration of VesselExpress analysis using increasing number of ROIs with an image dimension of 508 × 508 × 1000 μm. Every data point represents the doubling of previous image numbers starting with four. Analysis was performed either on a server (Intel® Xeon ® CPU E7-8890 v4, 2.20GHz, 96 cores, 1.97 TB RAM or a workstation (Intel® Xeon® W-1255 CPU, 3.30 GHz, 10 cores, 512 GB RAM).

**Fig. S2.**
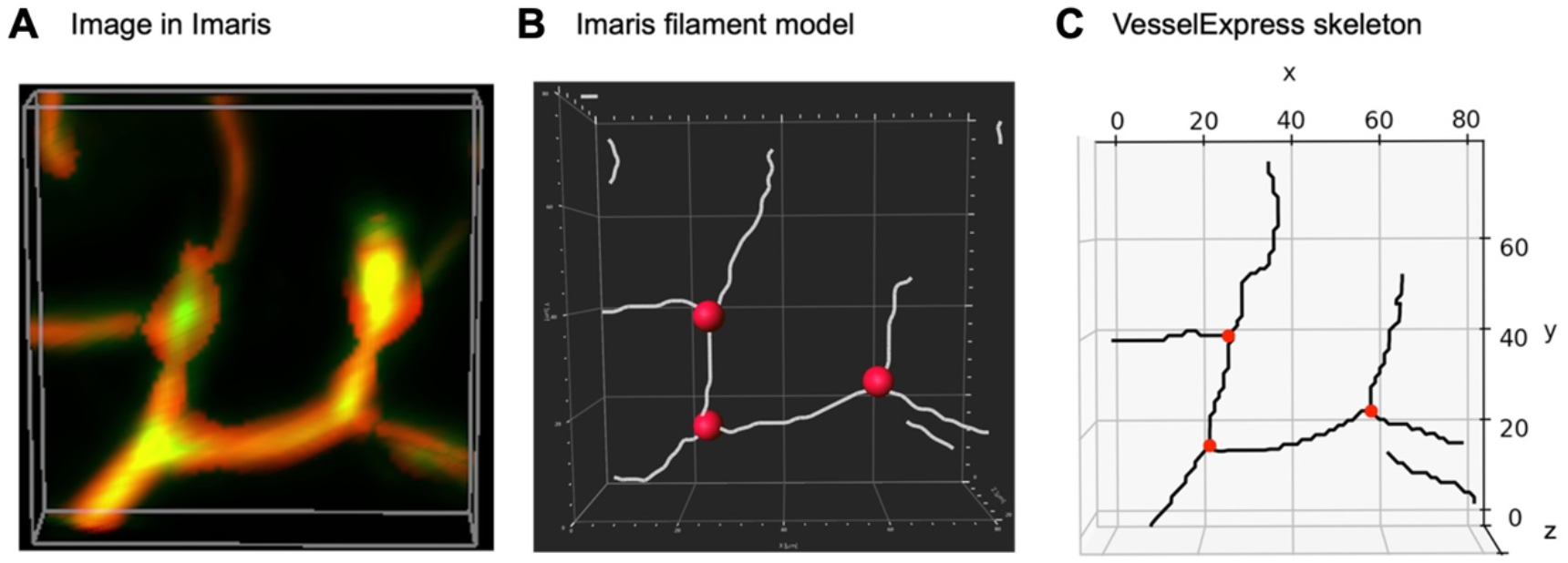
Visual validation of branching points. Branching points identified by VesselExpress were visually overlaid and compared with branching points identified by the Imaris based workflow. (A) Cropped region of a 3D LSFM mouse brain image visualized in Imaris. (B) Filament model obtained from the Imaris based workflow with branching points in red. (C) Skeletonization result of VesselExpress with branching points in red.

**Fig. S3.**
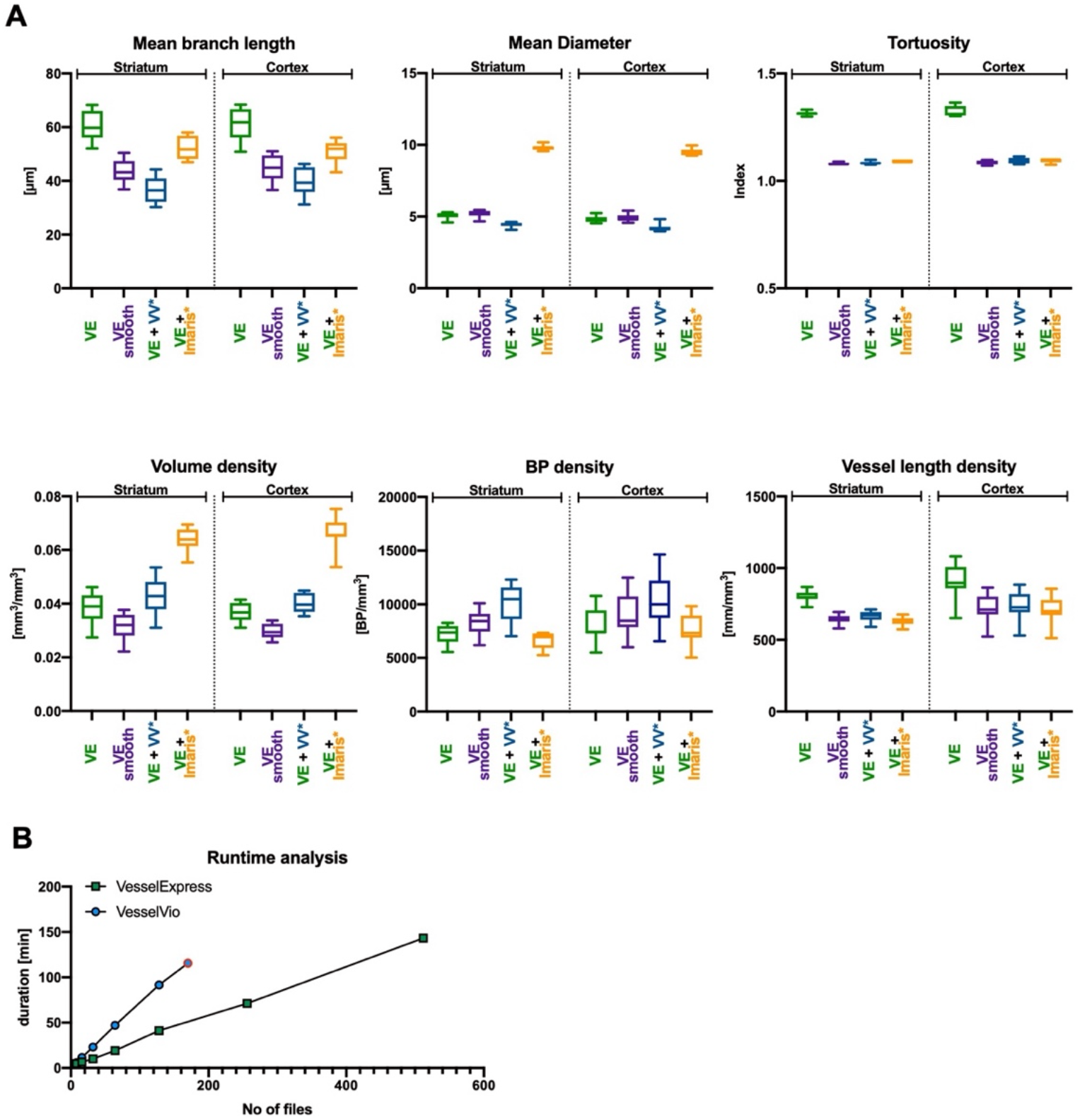
Comparison of VesselExpress with other analysis tools. (A) Analysis of blood vessels from brains of 12 weeks old healthy male mice in the striatum and cortex using VesselExpress (VE), VesselVio (VV) and Imaris shows that smoothing of Vessels results in 17.4% ± 1.3% smaller Vessel length density in VesselVio compared to VesselExpress. Furthermore, VesselVio detects considerably more branching points (38.5% ± 9.6%), resulting in shorter mean branch lengths (39.8% ±3.4%). The switchable smoothing step in VesselExpress (VE smooth) leads to comparable results as Imaris regarding vessel length and branching point density. Anyway, Imaris measures larger diameters and therefore a higher volume density. (B) runtime analysis between VesselExpress (without segmentation step) and VesselVio shows that VesselExpress is substantially faster than VesselVio. It is important to mention that the VesselVio encountered severe stability problems when dealing with more than 170 files (indicated with a red circle). A direct comparison with VesSap was not possible because the pretrained deep learning model provided by the VesSap authors yielded biologically wrong segmentations that were not usable for further downstream analysis. **\* Neither Imaris nor VesselVio provide segmentation methods for vessel segmentation, so that the segmentation provided by VesselExpress needed to be used to make comparison possible at all.**

**Fig. S4.**
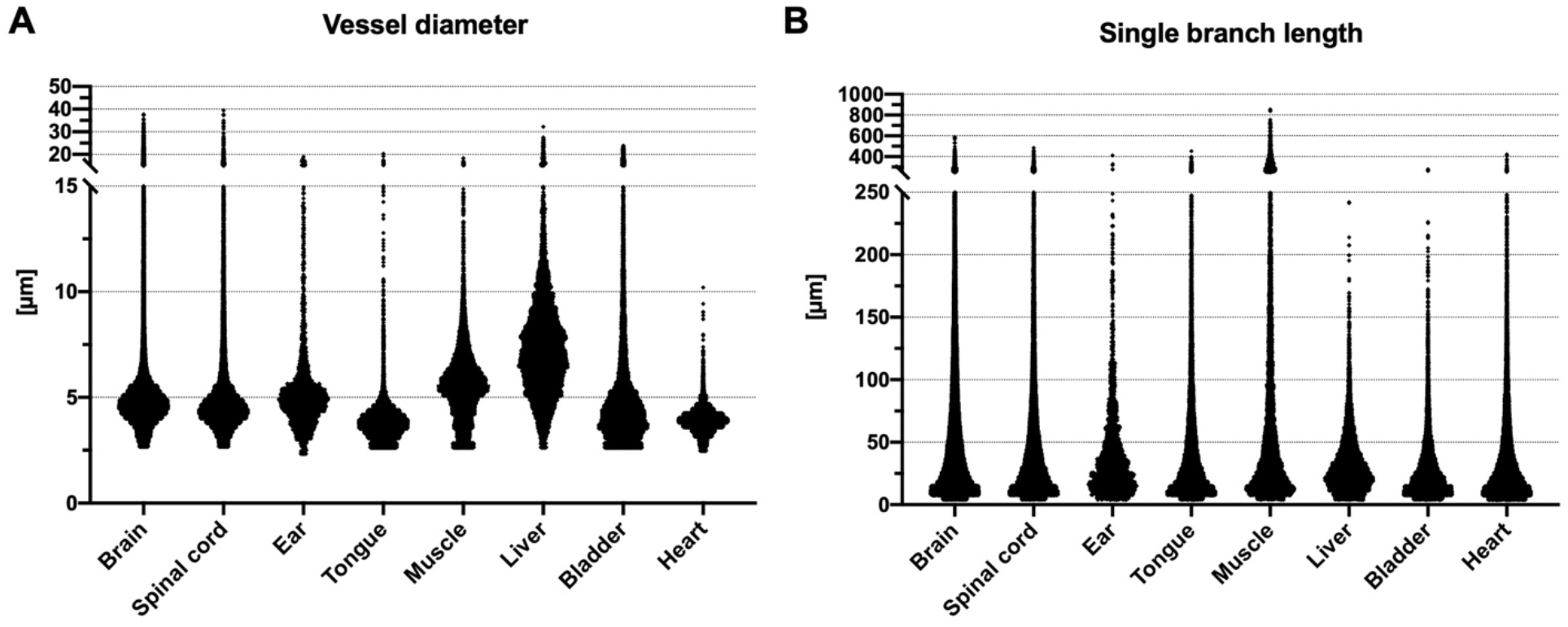
Dot plots representing single diameters (A) or vessel length (B) of all individual vessels in the respective organs.

**Fig. S5.**
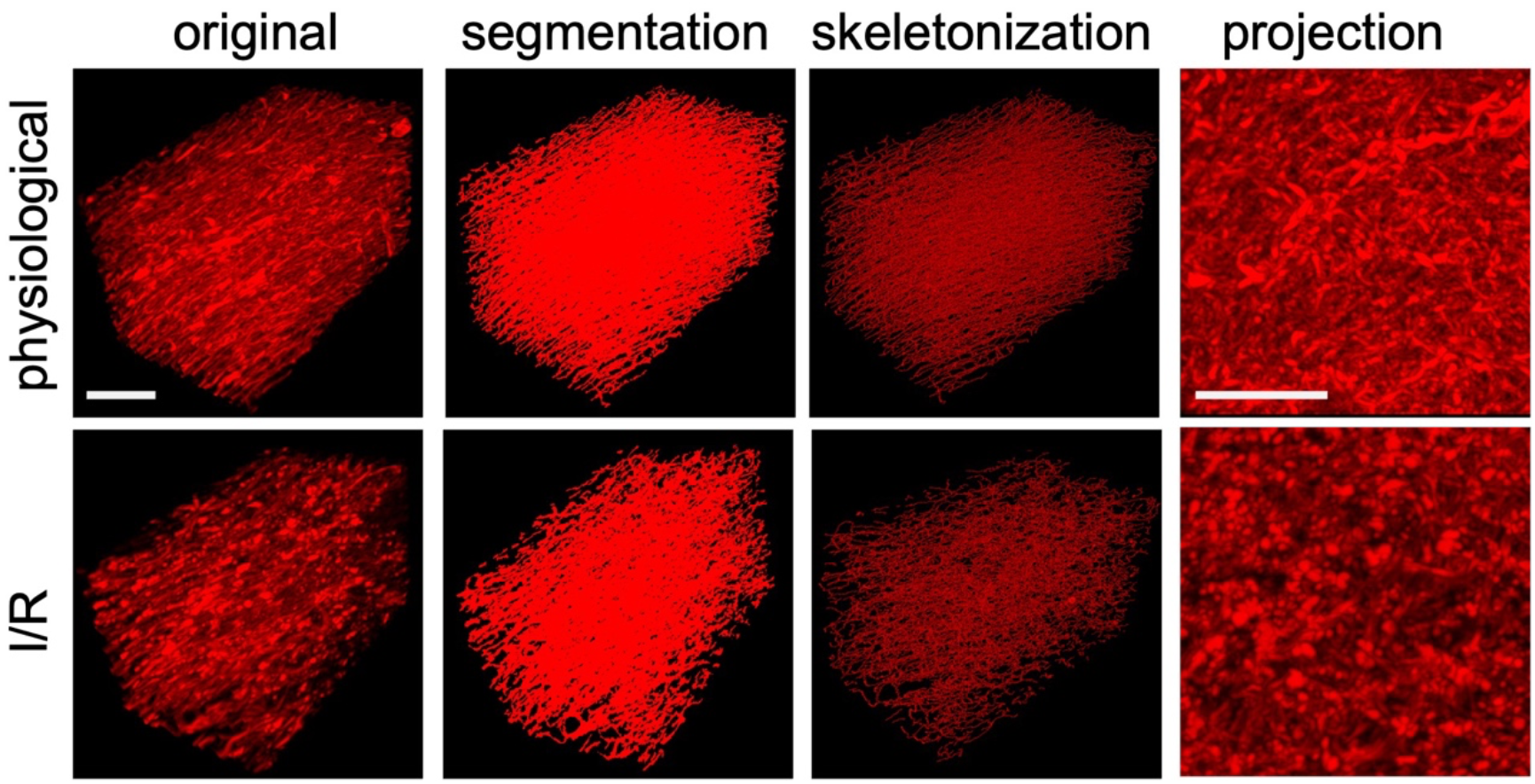
Representative images of healthy heart tissue (upper row) or infarcted heart tissue after 5 days of reperfusion (I/R; lower row). 2- or 3 Dimensional projections of original data as well as frangi conversion and skeletonized images obtained from VesselExpress analysis are shown. Scale bars represent 100 μm.

**Fig. S6.**
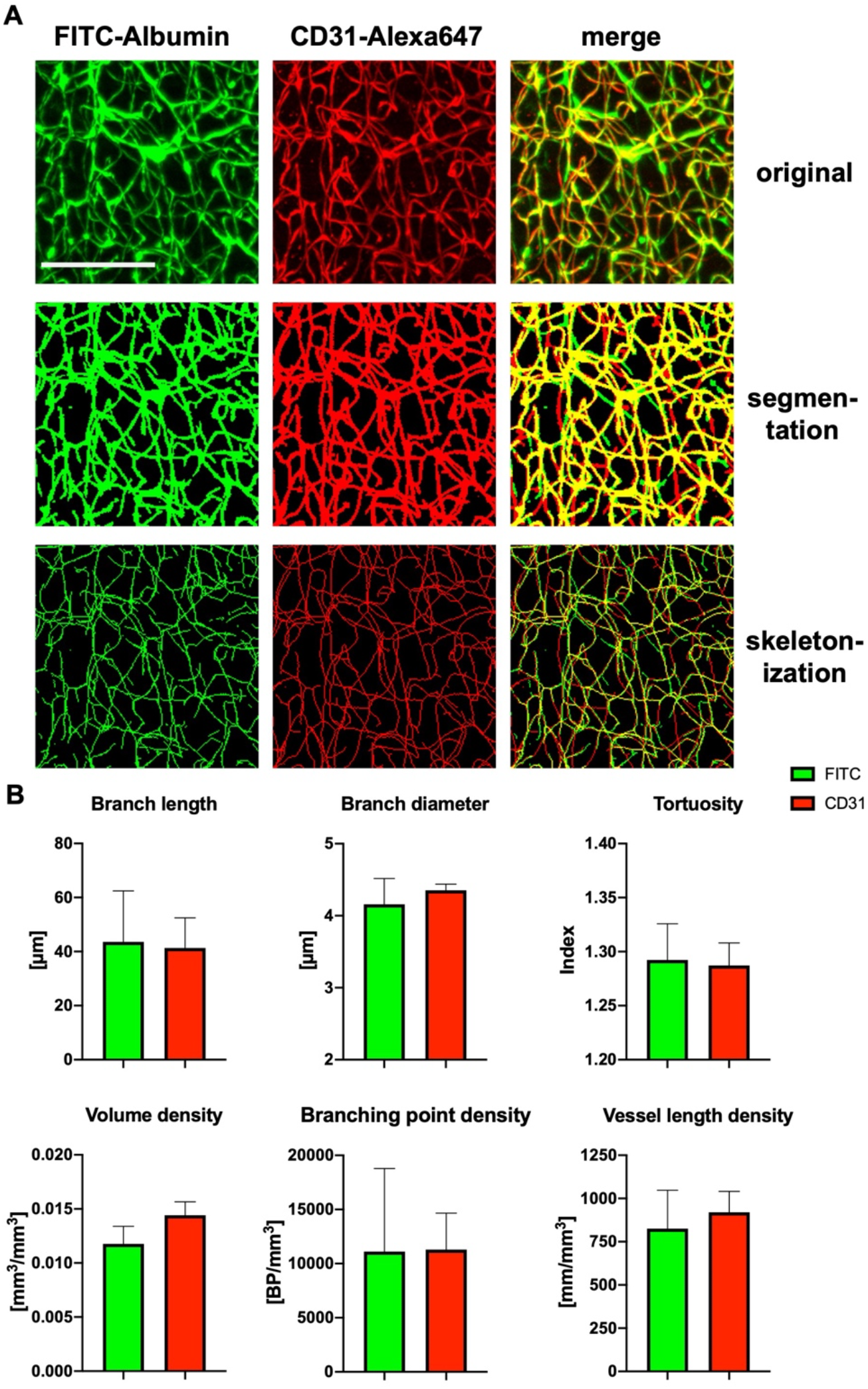
VesselExpress analysis of striatal vessels from brains of 12 weeks old healthy male mice labeled with FITC-albumin hydrogel and CD31-Alexa647 antibody. (A) original, segmented or skeletonized images as maximum projections in FITC-albumin-labeled brain vessels and the corresponding region labeled with CD31-Alexa647 antibody. (B) VesselExpress analysis of indicated parameters of 0.258 mm3 images of n = 4 mouse brain regions labeled with FITC-albumin or CD31-Alexa647 antibody, respectively. Scale bar represents 100 μm

**Fig. S7.**
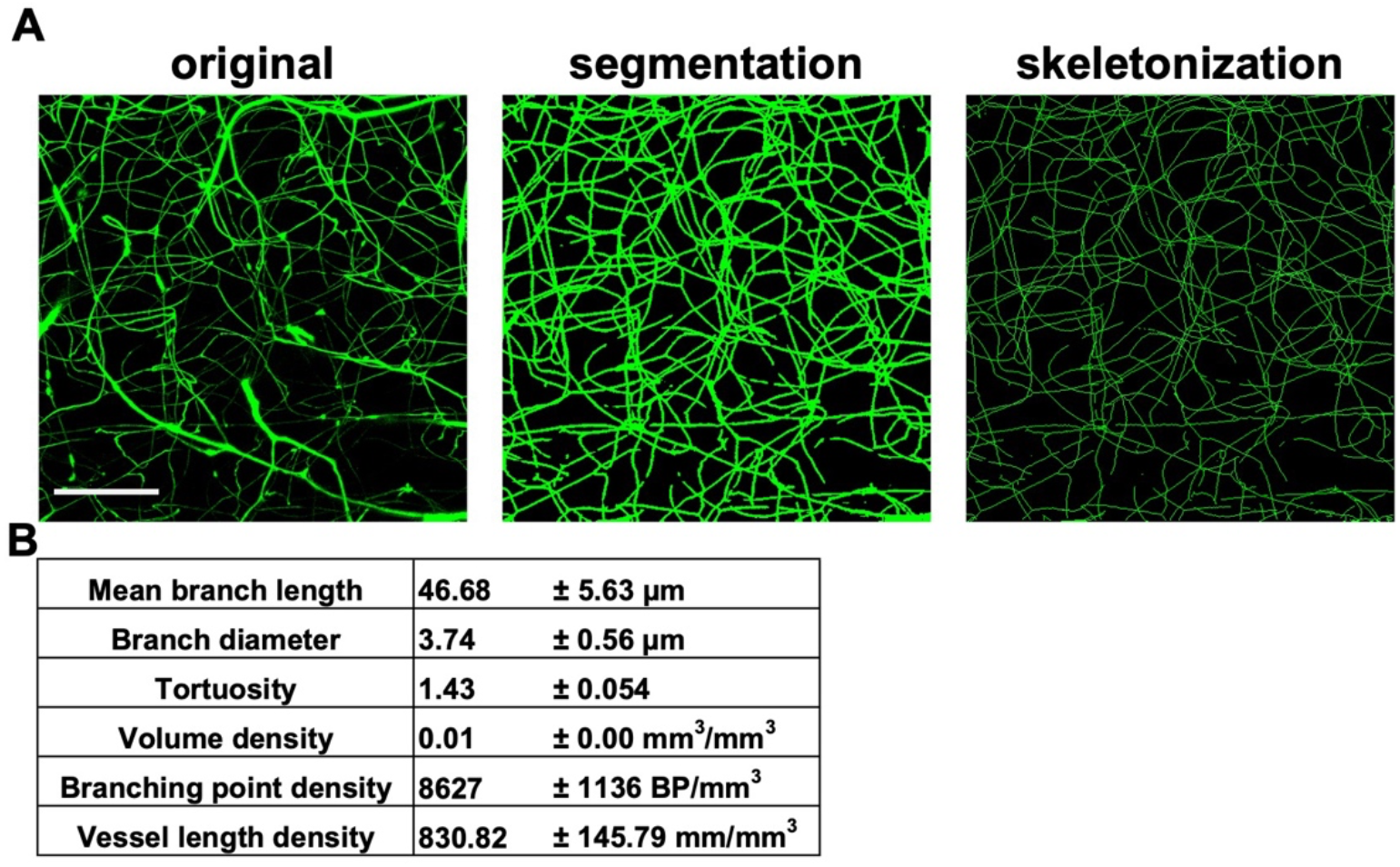
VesselExpress can be applied in images obtained with confocal microscope. (A) Maximum projections of confocal images of FITC-albumin-labeled striatal vessels from brains of 12 weeks old healthy male mice as original image, after segmentation and after skeletonization using VesselExpress. (B) Results of indicated parameters obtained using VesselExpress in confocal images (n = 4). Scale bar represents 100 μm.

**Table S1.** Image dimensions and image volumes of ROIs used for VesselExpress analysis.

| Organ | Image dimensions<br>(xyz [mm]) | Image volume<br>[mm<sup>3</sup>] |
| --- | --- | --- |
| Brain | 508 x 508 x 1000 | 0.258 |
| Spinal cord | 508 x 508 x 400 | 0.103 |
| Ear | 254 x 254 x 100 | 0.006 |
| Tongue | 305 x 305 x 200 | 0.017 |
| Muscle | 508 x 508 x 300 | 0.077 |
| Liver | 305 x 305 x 100 | 0.009 |
| Bladder | 305 x 305 x 200 | 0.019 |
| Heart | 253 x 253 x 555 | 0.036 |

## Supplementary Text

### Supplementary Methods

#### 1. Snakemake workflow

VesselExpress consists of multiple modules which are automated in a pipeline with the workflow management system Snakemake^16^. All modules of the pipeline are defined by rules in a Snakefile. A rule specifies input, output, environment, and the shell command to execute the corresponding Python script. Each module requires different software packages and contains parameters. The required software packages are defined in a YAML file. The default settings of all parameters are stored in a JSON configuration file which is also specified in the Snakefile. Thus, all parameters can be flexibly adjusted in the configuration file. Due to the modular software design, each processing step can also be executed individually. If preferred, custom functions can easily be integrated into VesselExpress by changing the rules in the Snakefile. All modules can be run by one terminal command (detailed instructions can be found on GitHub). Supplemental Figure S8 illustrates the workflow for processing a 3D image including all steps.

The workflow is also accessible through a web-browser based graphical user interface, as illustrated in Supplemental Video 1 which is linked on GitHub (https://github.com/RUB-Bioinf/VesselExpress). The video also provides guidance on how to adjust the workflow parameters.

**Fig. S8:**
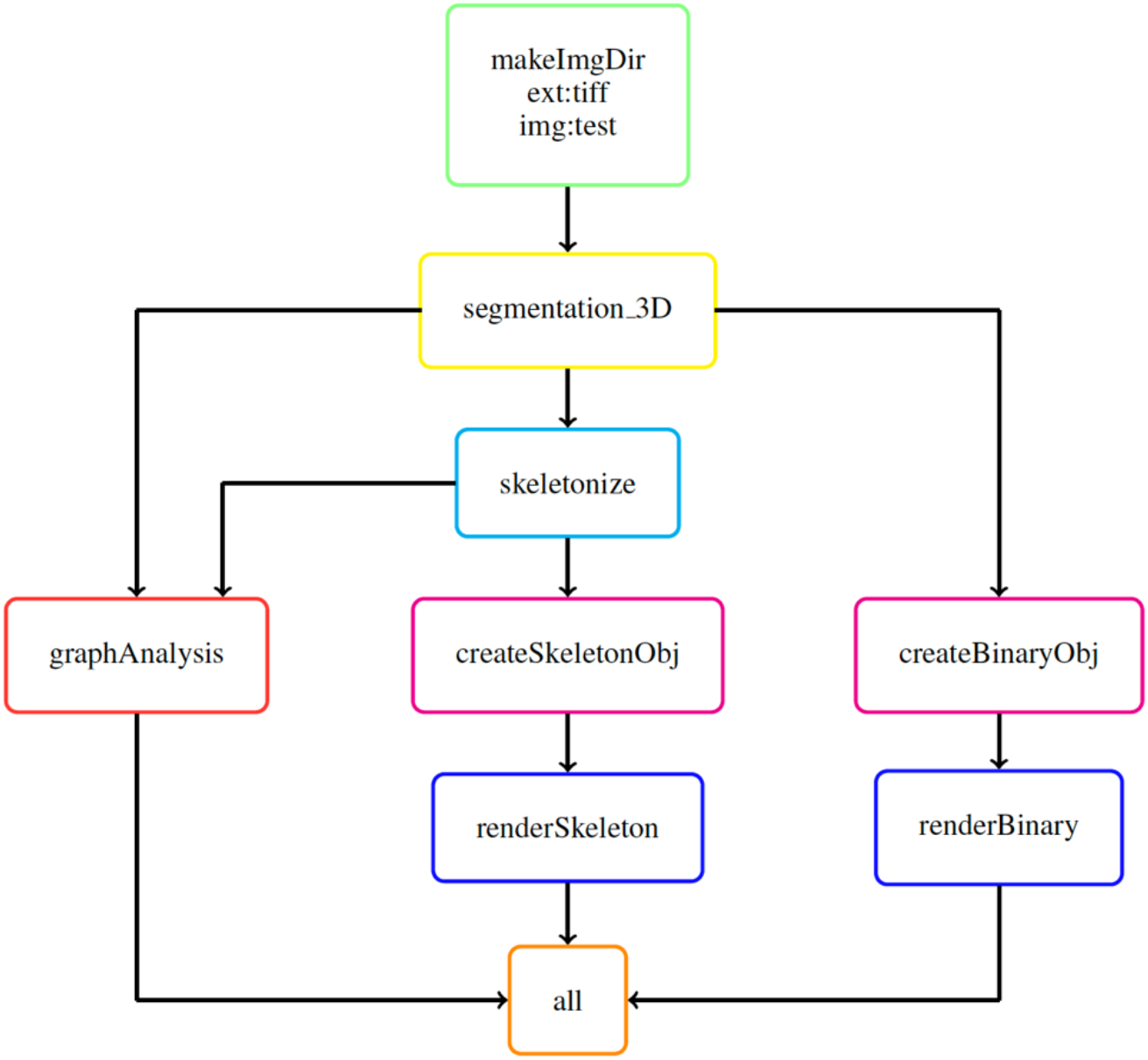
Visualization of Snakemake workflow for a 3D image as directed acyclic graph (DAG). Each node represents a rule of the Snakefile. First, a folder is created for the image to be processed (green) followed by the segmentation (yellow), which consists of 3 steps: pre-processing, core segmentation and post-processing. Next, the segmented image is skeletonized (light blue). This is followed by graph construction and analysis (red) which takes the binarized and skeletonized image as input. Optionally, the binarized and skeletonized images can be rendered. Therefore, the contours are first approximated via marching cubes (magenta) and then rendered in Blender (blue). In the rule, all (orange) output files are defined.

#### 2. Segmentation

The segmentation method is extended from classic image segmentation workflows from the Allen Cell and Structure Segmenter^17^, which is a 3-step workflow (pre-processing, core segmentation, post-processing) with a minimal number of parameters to tune and minimal number of functions to choose from. In the pre-processing step, we use the edge-preserving smoothing function in Segmenter. In the core segmentation step, we use a combination of a statistics-based thresholding method and customized Frangi filter-based segmentation. The statistical based thresholding method calculates the mean and standard deviation of the intensity of the whole image, denoted by *m* and *std*, respectively. The threshold value is set as *m + S × std*, where *S* is an empirically determined parameter (usually between 2.0 and 4.0, 3.0 was used for all organs). The customized Frangi filter based segmentation method takes three parameters: “sigma” (i.e., roughly representing the thickness of the vessels), “gamma” (i.e., the sensitivity to structures), and “cutoff method” (i.e., how to binarize the filter output into segmentation, options include Otsu, Li^3^, and Triangle^39^). Different parameters are optimized and used for different organs. Users can either load our pre-defined parameters for different organs or further adjust these parameters within a Napari plugin.

The statistics-based thresholding is mainly used to capture those very thick vessels with much higher intensity than the rest (e.g., main artery), while the rest (e.g., microvessels) will be picked up by the customized Frangi filter based segmentation. In certain organs, the variation of the thickness of different microvessels is too high for one single Frangi filter to segment accurately all vessel calibers (recall that the parameter “sigma” needs to be set according to vessel thickness). To address this issue, we used an innovative strategy employing two different Frangi filters with different parameters (optimized using the Napari plugin) to obtain accurate segmentation of various microvessels. It is worth mentioning that this “2 Frangi filters” method is different from the original multi-scale Frangi filter, where multiple sigma values are used in one single filter. For example, we were able to use one Frangi filter with sigma = 1, gamma = 5, cutoff method = Li and another Frangi filter with sigma = 2, gamma = 100, cutoff method = Otsu. However, using the original multi-scale Frangi filter with two sigma values (e.g., sigma = 1, 2) is only comparable to our “2 Frangi filter” method when both Frangi filters have the same gamma value and the same cutoff method. Without the flexibility of using different gamma values and different cutoff methods in the two Frangi filters, the original multi-scale Frangi filter usually suffers from over-segmentation errors (i.e., a thin vessel segmented much thicker than it should be or two proximal thin vessels falsely segmented as a merged one). The “2 Frangi filters” strategy is the key ingredient for successfully segmenting all vessels with accurate thickness in different tissues.

Before post-processing, a logical OR operation is applied to the output from the statistical based thresholding and the customized Frangi filter-based segmentations to combine the results. At last, the post-processing step ensues with three different functions to choose from, depending on the specific organ and imaging quality. These functions include topology-preserving thinning^12^ to refine the thickness of the segmentation, morphological closing to connect potentially fragmented vessels and small holes, and a size filter to remove small segmented objects due to noise or imaging artifacts.

#### 3. Skeletonization and graph construction

The vessels’ centerlines are extracted from the binary mask through the parallel thinning algorithm^40^ implemented in the scikit-image Python package^15^. The centerlines are then transformed into undirected graphs by using the Python 3scan toolkit^20^ (Supplemental Figure S9). The graph construction consists of two main steps: first vertices are created for each image point (Supplemental Figure S9A and S9B). Vertices are defined by their image coordinates. Neighboring vertices are connected via edges. After this step, the graph typically contains cliques, i.e., subgraphs in which all vertices are connected, resulting in an over quantification of branching points and underdetection of terminal points (vertices with one neighbor). To avoid this, all cliques comprising three vertices are resolved by removing the longest edge (Supplemental Figure S9C). The final graph is stored as an adjacency list. For each vertex, there is an adjacency list that contains all neighboring vertices.

**Fig. S9:**
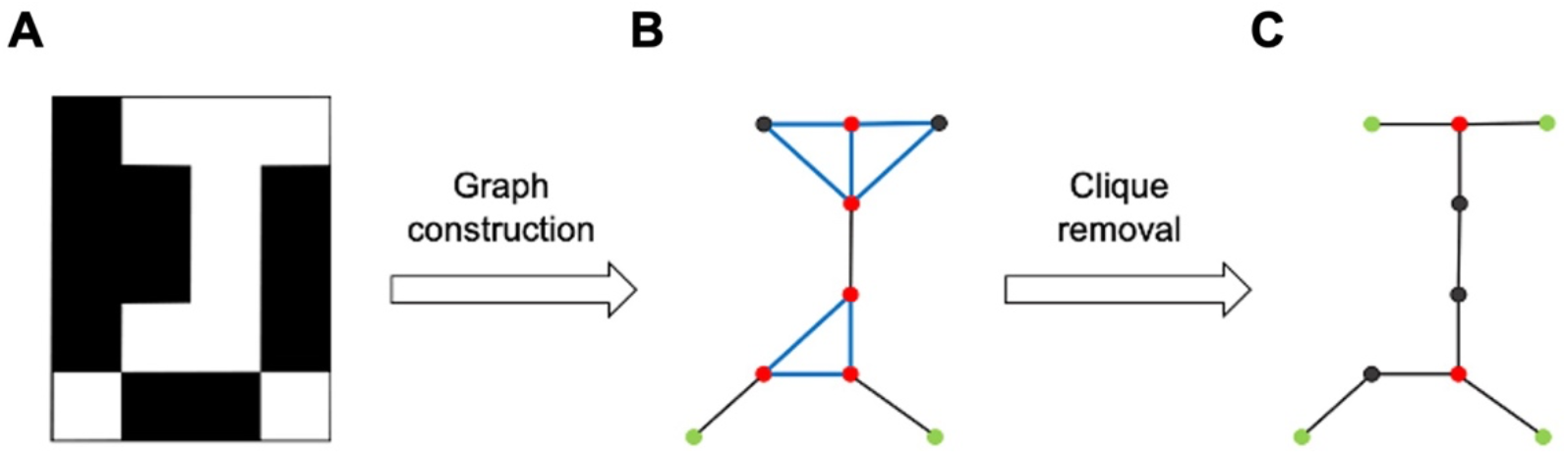
Graph construction from a skeletonized binary image. (A) Binary skeleton image with white foreground and black background. (B) Each foreground point is represented by a node in the graph. All neighboring nodes are connected via edges. This creates cliques (blue) and thus too many branching points (red) and not enough terminal points (green). (C) After removing cliques, the graph contains the correct number of branching and terminal points.

The graph obtained from skeletonization may contain spurious branches that do not represent the topology of the object. These occur when the binary image has bumps on the edge of the object. Therefore, these are branches that lie at the edge of the object and thus start from the center of the object in a branching point and end in a terminal point at the border of the object. These branches are identified and removed by the criteria defined by Montero and Lang ^41^ found in equation 1:

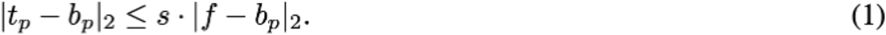

A branch with branching point *b_p_* and terminal point *t_p_* is removed exactly when the length of a branch is less than or equal to the distance of the branching point *B_p_* to the nearest background point *f* (Supplemental Figure S10). With the scaling factor *s,* this distance can be scaled arbitrarily. The higher the scaling factor, the more branches are removed (“pruned”). The nearest background point *f* is determined using the Euclidean distance transformation from the scikit-image Python package ^42^.

**Fig. S10:**
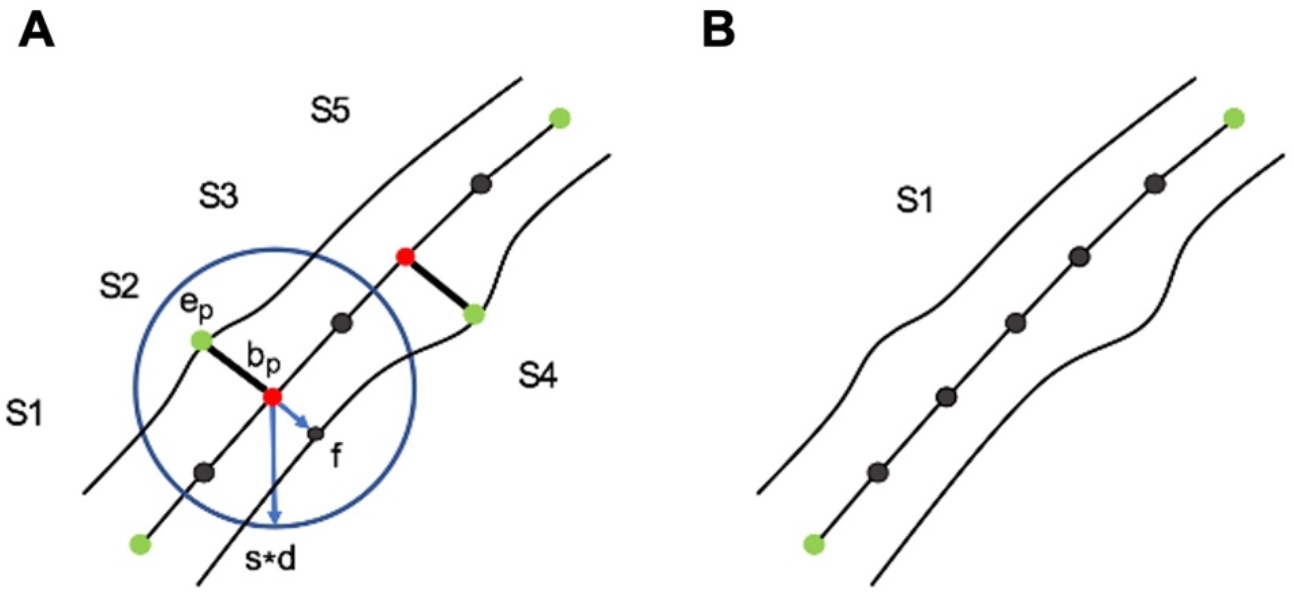
Pruning of spurious branches. (A) Branches with a length smaller than the distance of the branching point *b_p_* to the nearest background point *f* multiplied by *s* are removed. (B) The graph consists of only one segment after pruning.

#### 4. Graph analysis

The graph is used to extract relevant features of the vascular network. The features can be divided into filament and segment statistics. Each connected component of the graph is identified as one filament, and each direct path between branching or terminal points is considered as a segment. Supplemental Figure S11 shows a connected graph, which consists of five segments. To identify segments, each connected graph is traversed in a depth-first search (DFS) starting from a randomly chosen terminal point. For each vertex, the number of neighbors is read out from the adjacency list to check if it is a branching or terminal point. Once a branching or terminal point is found, the segment is backtracked to the last terminal or branching point. If a previously visited vertex is found, it is designated a circle. Once a segment is found, all features of the segment are calculated.

In a post-processing step edge points which are located at the border of an image are excluded from the terminal point quantification since they cannot be considered actual terminal points. Also segments with length or diameter below user-defined thresholds are removed from the analysis.

For each filament, the number of branching and terminal points is counted. In addition, the number of neighbors of each branching point is read out from the adjacency list. The total vessel length is calculated from the sum of all segment lengths. The length of a segment corresponds to the sum of the Euclidean distance of all points from starting point *i* to terminal point *e* of the segment *S* according to the pixel dimensions *d*:

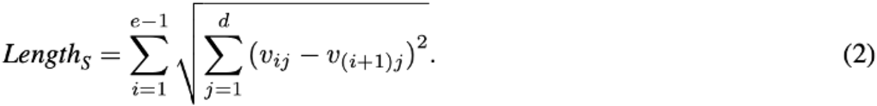

**Fig. S11:**
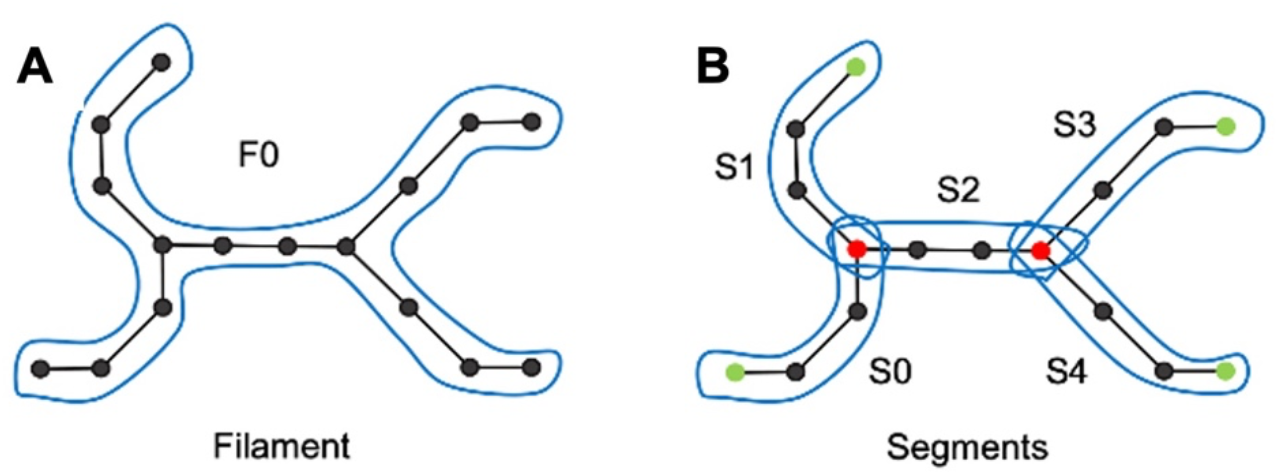
Filament vs segments. (A) A connected graph is called a filament (marked F0). (B) The graph consists of 5 segments (numbered S0-S4). A segment is a branch between branching/terminal points.

Furthermore, the tortuosity is calculated for each segment with the equation 4. Equation 3 is used to calculate the Euclidean distance between the starting point *s* and the terminal point *t* for each dimension *d* of a segment. VesselExpress outputs the quotient of the Euclidean distance *h_S_* and *lenhth_S_*.

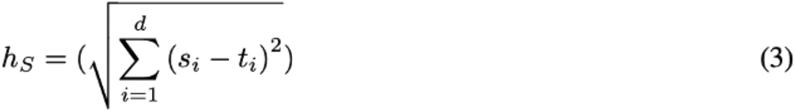

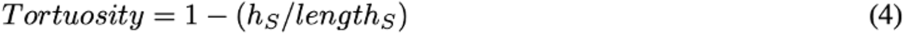

In addition, diameter and volume of the vessels are determined. The decisive value is the radius. This is calculated by the Euclidean distance transformation using equation 5. The distance transformation is carried out on the binary image and replaces the pixel values *x* with the Euclidean distance of the point to the nearest background point *b* considering the *n* dimensions. The diameter results from the average of all radii of the *m* skeleton points of a segment from equation 6.

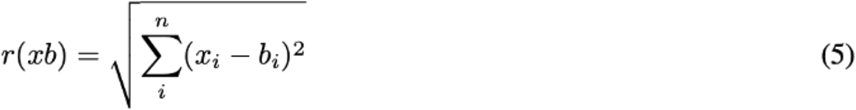

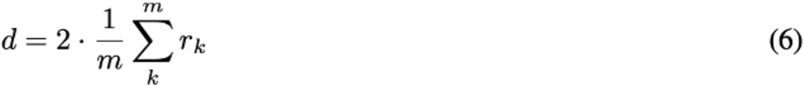

The vessel volume is calculated by using the vessel’s average radius:

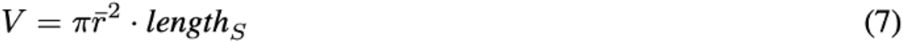

For segments that are connected to a predecessor via a branching point, branching angles are also determined. The angles are calculated with the standard angle equation for two vectors:

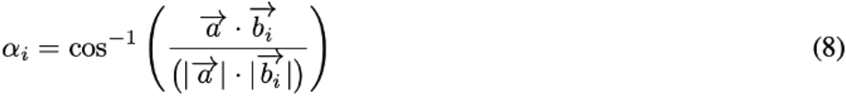

where *i* corresponds to the considered segment and *b_i_* corresponds to the respective adjacent segment of the predecessor *a*. A threshold value specifies the percentage length of the segment, from which the vector of a segment is formed.

Therefore, the value range of the threshold is between 0 and 1. If 0, the first neighboring node and if 1, the last node is selected as an approximation of the vessel orientation (Supplemental Figure S12).

**Fig. S12:**
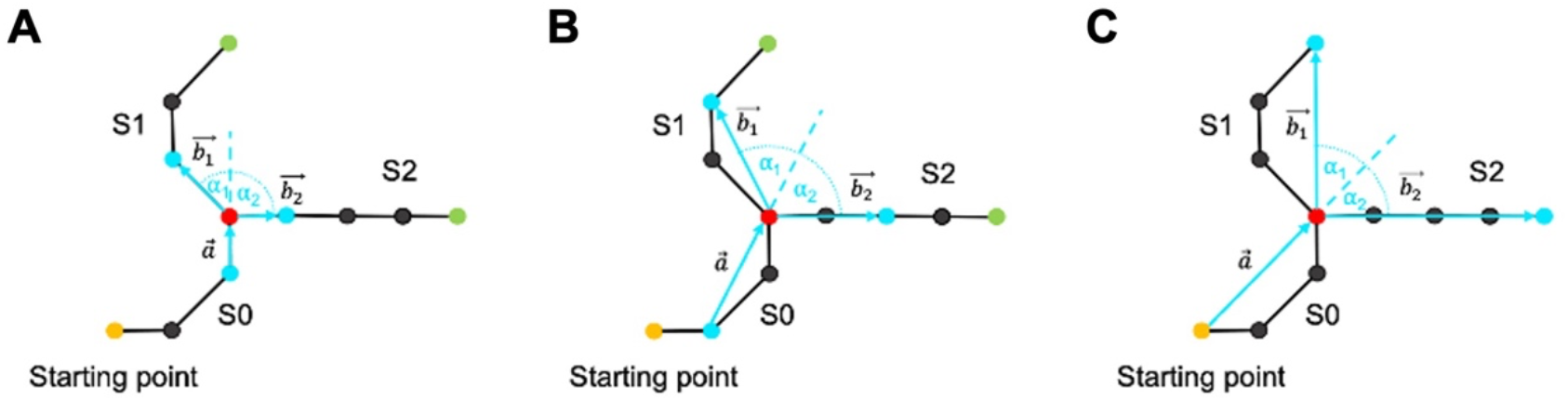
Effect of different threshold values on the branching angles. The predecessor segments are determined starting from the starting point (yellow). The threshold value indicates the length from the branching point (red) to the point from which the vector is formed (blue). (A) Threshold = 0. (B) Threshold = 0.5. (C) Threshold = 1.

#### 5. Rendering

VesselExpress features an automated Blender^21^ integration, running on Version 2.83.5. Using the provided Cycles or Eevee rendering engines, VesselExpress can automatically generate high-definition images of the calculated segmentation as well as the skeleton.

This rendering makes full use of the Blender capabilities using the Principled BSDF material for the meshes and raytraced rendering on the GPU (if available). The mesh-, material-, lighting-, and color-properties, as well as camera angles can be freely customized beforehand.

The meshes for rendering are created via the scikit-image implementation of the Marching Cubes Algorithm ^43,44^. As an optional output, the generated meshes can be saved for further use in form of a .stl, .glb, and .blend file each.

#### 6. Validation on synthetic ground truth dataset

To evaluate the accuracy against absolute ground truth, we created a synthetic dataset with each image containing six tubes of a certain size and diameter. The images were generated to replicate the resolution of the original light-sheet microscopic images as closely as possible. For each image, we compared the length and diameter of the pipeline results (actual value) of VesselExpress or VesselVio with the expected target values. Supplemental Table S2 shows the results. This comparison reveals that lengths are precisely calculated (average deviation of 1.58% or 0.41%) as well as vessel diameters (average deviation of 5.61% or 3.04%) using VesselExpress or VesselVio, respectively.

**Table S2:** Validation of VesselExpress and VesselVio against a synthetic ground truth

| Image index | Vessel Length |  |  |  |  | Vessel diameter |  |  |  |  |
| --- | --- | --- | --- | --- | --- | --- | --- | --- | --- | --- |
|  | Target (µm) | VesselExpress |  | VesselVio |  | Target (µm) | VesselExpress |  | VesselVio |  |
|  |  | Actual (µm) | Deviation (%) | Actual (µm) | Deviation (%) |  | Actual (µm) | Deviation (%) | Actual (µm) | Deviation (%) |
| 1 | 125 | 124.00 | -0.80 | 122.00 | -2.40 | 3 | 4.00 | 33.33 | 4.43 | 47.59 |
| 2 | 250 | 248.98 | -0.41 | 250.92 | 0.37 | 6 | 4.55 | -24.17 | 5.18 | -13.67 |
| 3 | 375 | 368.33 | -1.78 | 371.25 | -1.00 | 9 | 8.56 | -4.89 | 10.26 | 14.04 |
| 4 | 500 | 491.00 | -1.80 | 500.82 | 0.16 | 12 | 8.42 | -29.83 | 10.69 | -10.90 |
| 5 | 625 | 611.60 | -2.14 | 619.53 | -0.87 | 15 | 14.39 | -4.07 | 15.55 | 3.65 |
| 6 | 750 | 738.00 | -1.60 | 752.76 | 0.37 | 18 | 17.89 | -0.61 | 18.10 | 0.55 |
| 7 | 875 | 857.00 | -2.06 | 867.82 | -0.82 | 21 | 20.26 | -3.52 | 21.12 | 0.59 |
| 8 | 1000 | 985.00 | -1.50 | 1004.70 | 0.47 | 24 | 20.20 | -15.83 | 20.35 | -15.22 |
| 9 | 1125 | 1100.80 | -2.15 | 1113.30 | -1.04 | 27 | 26.04 | -3.56 | 28.68 | 6.23 |
| 10 | 1250 | 1230.00 | -1.60 | 1258.68 | 0.69 | 30 | 29.12 | -2.93 | 29.27 | -2.44 |

#### 7. Confocal image acquisition

For confocal microscopy, 1 mm brain sections dehydrated and cleared by the CUBIC method ^45^ were used. For this purpose, PFA-fixed brains from mice were incubated in CUBIC-1 reagent for 5 days with constant shaking while the solution was changed daily resulting in completely clear tissue. Subsequently, the image stacks from cubic cleared brain slices were acquired on a Leica TCS SP8 inverted confocal microscope equipped with a white light laser source combined with an acousto-optic tuneable filter for excitation wavelength selection and spectral detection on hybrid photodetectors via an acousto-optic beamsplitter. For image stack acquisition, an HCPL Fluotor 10X/0.3 air objective was used and the confocal scanner configured to acquire 266 image frames of 512 × 512 pixels (0.994 micron pixel dimension xy) in the z-direction with a z-stepping size of 4 microns. The FITC-albumin hydrogel filled brain vasculature was excited with a wavelength of 488 nm and the fluorescence emission detected from 500 nm to 580 nm.

#### Movie S1

https://github.com/RUB-Bioinf/VesselExpress

## Notes

https://zenodo.org/record/6025935#.YyGN5S0RqJ8

### Competing Interest Statement

The authors have declared no competing interest.

